# Finite Element Modeling of the Scaphoid Shift Maneuver: Implications for Scapholunate Ligament injuries

**DOI:** 10.64898/2026.02.17.705556

**Authors:** Thor E. Andreassen, Taylor P. Trentadue, Andrew R. Thoreson, Parunyu Vilai, Kai-Nan An, Sanjeev Kakar, Kristin D. Zhao

## Abstract

**Background:** Computational modeling is a tool being deployed for orthopaedic solutions but its use in the hand and wrist remains limited. This work used a model to simulate a clinically relevant provocative scaphoid shift maneuver (SSM) with different levels of scapholunate interosseous ligament (SLIL) injuries to observe the effect on different metrics.

**Methods:** A personalized model simulated the full SSM motion cycle from ulnar deviation with extension to radial deviation with flexion informed by the participant’s motion obtained from dynamic computed tomography. Models repeated the SSM under different levels of SLIL injury and reported changes in joint kinematics, contact mechanics, and ligament forces.

**Results:** The fully injured model increased scaphoid dorsal translation, flexion, and radial deviation compared to the intact condition and caused a subluxation of the scaphoid. Radioscaphoid contact areas were approximately 200% greater in the fully injured model compared with all others and the fully injured model was the only condition where contact force decreased across the motion cycle. Ligament forces in the intact condition were on average 33.0 N and 54.2 N for the volar and dorsal SLIL, respectively. Lastly, the long radiolunate, an extrinsic stabilizer, had forces that increased following SLIL injury.

**Conclusions:** Computational models can successfully recreate clinically observed behaviors of an SSM, including scaphoid subluxation, while providing new insights via quantification of contact mechanics and ligament forces. Contact mechanics metrics may be important for understanding the long-term progression of untreated SLIL injuries to osteoarthritis. Additionally, ligament force metrics may explain the progression of SLIL injuries from volar SLIL to dorsal SLIL and highlight the importance of repairing extrinsic stabilizers of the joint, due to increased force sharing following SLIL injury. This work provides a pathway to future studies investigating the effects of SLIL injury and repair, both acutely and chronically.

## 1 Introduction

Computational modeling in biomechanics remains a fundamental tool in research^1^ and industry settings^2^. The growing importance of personalized medicine^3^, along with recent trends in digital twins^4-6^ and *in silico* clinical trials^7^, has led to an increased prominence of these models in clinical-translational orthopaedics to predict surgical outcomes^8^, evaluate new joint arthroplasty component designs^9^, and patient-specific implant assessment^10^. Still, compared to the lower extremity, the use of computational models in the upper extremity remains limited.

Of the six major appendicular joints (ankle, knee, hip, wrist, elbow, shoulder), finite element modeling (FEM, a computational modeling technique frequently used in engineering) has been least frequently applied to the wrist^11^. Wrist modeling remains difficult because of the large number of structures (bones, ligament, cartilage, etc.) and a paucity of open-source experimental data^12^. Moreover, most models are limited to static or simplified simulations of joint forces or motion^12,13^. As such, they do not adequately capture the true complexity of the wrist joint for clinical-translational applications. Enhancing the complexity and realism of these models will significantly increase their applicability in clinical-translation settings.

One potential use of FEM is recreating clinical exam maneuvers. For instance, the scaphoid shift maneuver (SSM, also called scaphoid shift test or Watson test)^14^, is often performed to identify patients with scapholunate interosseous ligament (SLIL) tears^15^. The SSM is frequently described as moving the wrist from full ulnar deviation with slight extension to full radial deviation with slight flexion while maintaining dorsal-directed pressure on the scaphoid. Positive tests include excessive dorsal motion or subluxation of the scaphoid^16^ and have approximately 45-80% sensitivity and 62-71% specificity^16-18^ for diagnosis of SLIL tears. These provocative maneuvers are crucial in clinical settings^19^, but have not been explored with computational modeling^12^.

While the exact ligaments injured depend on a range of factors including injury mechanism, severity, and patient-specific factors, SLIL tears commonly start at the volar SLIL (VSL), progressing to the proximal SLIL (PSL) and lastly the dorsal SLIL (DSL)^20,21^. Severe injuries increase coronal plane scaphoid-lunate gap (in extreme cases exhibiting scapholunate diastasis) and dorsally shift the scaphoid in the sagittal plane^20,22^. In cases of injury, the scaphoid and lunate bone rotate in opposition, with the scaphoid flexing, particularly with radial deviation^20^. The SSM is designed to accentuate these behaviors. Computation models can recreate these clinical patterns by quantifying the scaphoid’s kinematic changes in response to injury: specifically, dorsal-volar translation, radial-ulnar translation, and flexion-extension rotation. Moreover, these models may provide new insights by quantifying metrics that are otherwise challenging to observe, e.g. joint contact mechanics (forces, areas, and pressures) and ligament forces. These are impractical to measure with conventional methods but are a common output of computational methods.

Our goal was to use a validated model of the wrist, developed from a healthy participant dynamically imaged with four-dimensional computed tomography (4DCT = 3DCT + time), to simulate a SSM for an intact wrist and at three levels of SLIL injury. The models predicted bone kinematics, joint contact mechanics, and ligament forces during the SSM. We hypothesized that (1) the scaphoid will have the greatest dorsal translation, radial translation, and flexion in the model with a complete SLIL injury; (2) A complete SLIL injury will cause the most dramatic changes to the joint contact mechanics; and (3) there will be increased force sharing in the surrounding extrinsic ligaments with increased SLIL injury severity.

## 2 Materials and Methods

An overall workflow is provided in Figure 1. The models used were developed from an existing personalized wrist model from our prior work^11^ and driven, in part, using experimentally-obtained joint motion^23,24^. The novelty of this work includes methods to simulate an SSM with different types of SLIL injury to observe kinematics, ligament forces, and contact mechanics.

**Figure 1.**
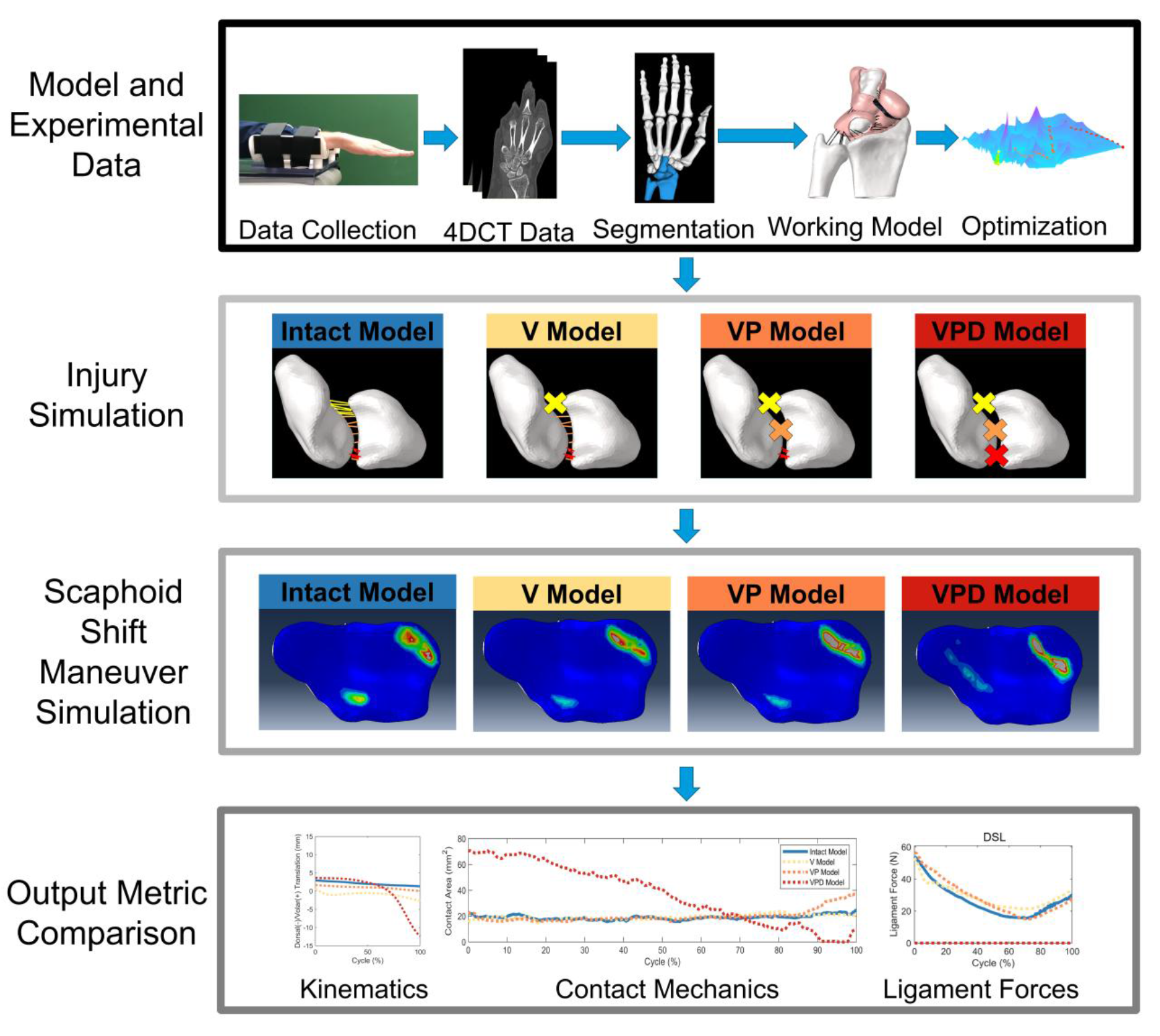
Overall study workflow. The portion in the black box represents original model and experimental data from prior work^11^. The gray boxes represent the methods employed to simulate injury, simulate the scaphoid shift maneuver, and then compare output metrics of interest including joint kinematics, joint contact mechanics (contact area, contact force, and contact pressure) and ligament forces. The four models built are the intact model (intact SLIL), V model (VSL injury), VP model (VSL and PSL injury), and the VPD model (VSL, PSL, and DSL injury).

**Figure 2.**
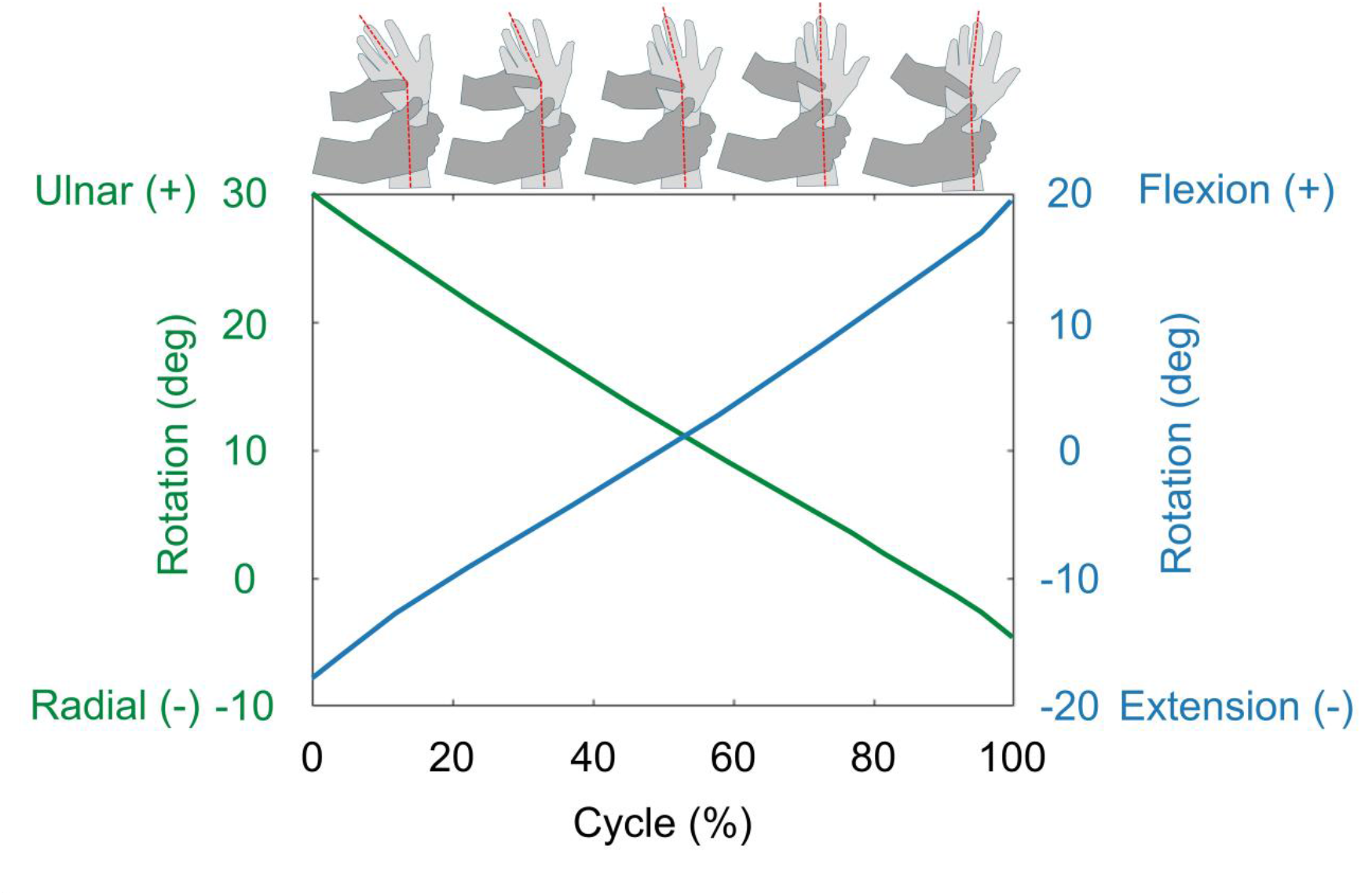
Simulated scaphoid shift maneuver (SSM) kinematics of the right hand of a participant. Range-of-motion bounds were empirically determined from 4DCT. Models driven with ulnar deviation and slight extension (cycle = 0%) to radial deviation and slight flexion (cycle = 100%) while maintaining a small constant force of 25 N dorsally on the tubercle of the scaphoid.

### 2.1 Model and Experimental Data

A personalized computational model was developed in Abaqus Explicit (Dassault Systemes, France) for an individual (Female, Age 39, Type I lunate-hamate facet) to expand upon previous work^11^ using data collected under an IRB-approved study(IRB 20-007668)^23,24^. The details of model development have been previously described^11^ but will be summarized herein. Briefly, participants were imaged in neutral wrist positions using static CT and during a range of dynamic tasks using 4DCT. Specifications of the data acquisition methods and parameters have been published previously^25,26^. Individual bones were segmented from static CT volumes using Analyze (Mayo Foundation for Medical Education and Research, Rochester, MN) and used to create 3D surfaces. Our existing registration pipeline was used to determine the position of the bones in each of the volumes of the 4DCT^27,28^. In the computational model, the bones were modeled as rigid structures.

A novel pipeline, based on a publicly-available non-linear morphing algorithm^29^, was applied to the bones to automatically predict the location of cartilage and ligament attachment sites. This enables prediction of soft tissue attachment sites without manual segmentation of soft tissue structures by mapping the positions identified on a template anatomy to each new instance^11^.

Using the automated soft tissue sites, a combination of algorithmic techniques—including surface projection (extracts cartilage thicknesses), *k* means (identifies ligament fiber endpoints), and the Kuhn-Munkres algorithm (matches ligament endpoints to create fibers) —were used to create the cartilage structures and individual ligament fibers^11^. Cartilage was modeled as rigid but with an optimized non-linear contact pressure-overclosure relationships between interfaces^30^, and ligaments were modeled as non-linear tension only springs^30^. The completed structures were verified for model compatibility in HyperMesh (Altair Engineering Inc., Troy, MI). The model encompasses the radius, capitate, scaphoid, lunate, and ulna bones; cartilages of the distal radius, capitate, scaphoid, and lunate; as well as representations of the volar SLIL (VSL), proximal SLIL (PSL), dorsal SLIL (DSL), long radiolunate (LRL), short radiolunate (SRL), radial collateral (RCL), radioscaphocapitate (RSC), ulnolunate (UL), dorsal radiocarpal (DRC), scaphocapitate (SCL), and ulnocapitate (UCL) ligaments.

The resulting model applied the participant’s experimentally-obtained kinematics during an unresisted radial-ulnar deviation task. Ligament material properties—including stiffness and reference strain or resting length—were optimized to minimize differences between experimentally-obtained kinematics and model-predicted kinematics, adapting a pipeline previously deployed in the knee^30,31^. Per recent recommendations for model credibility^32^, the model was validated by predicting the scaphoid kinematics in an unseen joint motion, namely a radial-ulnar deviation motion against resistance. The model predicted scaphoid kinematics were close to the experimentally-obtained motion with translations within 0.75 mm and rotations within 3.75°.

Ultimately, this yielded a model with the participant’s unique osseous geometries, individual predicted cartilage and ligament insertions, and with ligament properties optimized to recreate their experimentally-obtained joint motion. This model was then used to simulate the SSM.

### 2.2 Scaphoid Shift Maneuver (SSM) Simulation

The model sought to recreate the effects of a clinical SSM by applying relevant dynamics to the modeled carpal bones. The model applied wrist movement using displacement control for the capitate and force control for the scaphoid and lunate. This was achieved through a simulated SSM involving ulnar deviation with extension (approximately 30 degrees of ulnar deviation and 20 degrees of extension at 0% motion cycle) through radial deviation with flexion (approximately 10 degrees of radial deviation and 20 degrees of flexion at 100% motion cycle). This approach is based on the motion of the capitate, as it closely approximates the overall position of the hand relative to the forearm^33,34^. The range of radial-ulnar deviation was taken from the participant’s obtained range of motion during a deviation task, and the extension-flexion range was derived from prior work^35,36^. All kinematics reported used a radial-based coordinate system that was applied to the volumetric centroid of all other bones in the static CT pose.

The dorsal force applied during an SSM was included in the model as a small constant force of 25 N, based on the approximate average force in prior work^37^, directed dorsally at the volumetric centroid of the scaphoid. Whereas true application of this force would be at the scaphoid tubercle, representing bones as rigid bodies within the model provides an opportunity to simplify the definition of the force with minimal consequence by applying the force at the volumetric centroid (Figure 3). The small moment arm of 2.7 mm in the worst case (Figure 3) means that a minimal flexion-extension moment of 0.068 N*m would be neglected with this approach. As such, this simplification is valid with a minimal impact on the dynamics involved.

**Figure 3.**
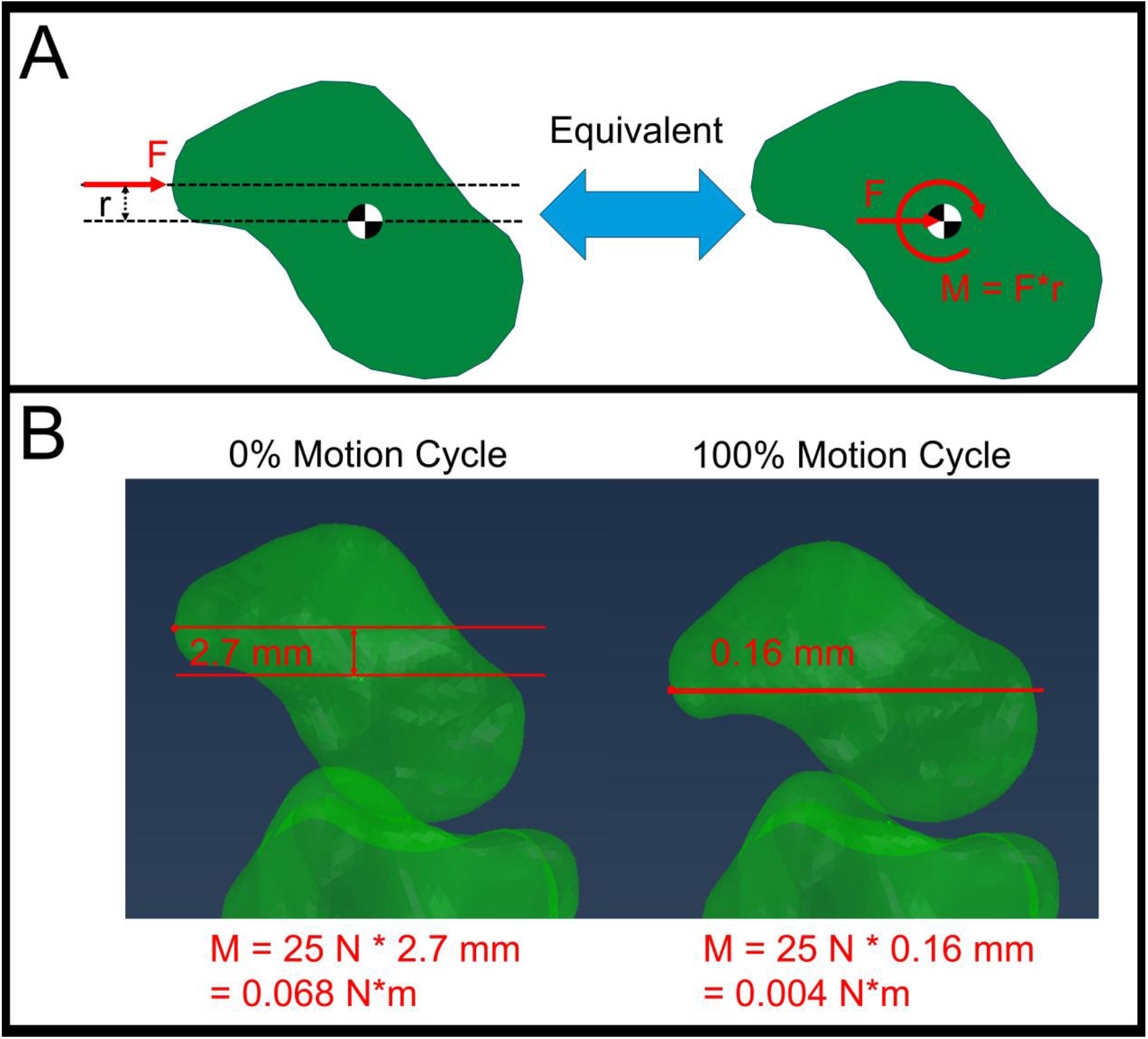
(A) Free body diagram of the scaphoid showing the moment that is created by the force applied to the distal tubercle vs. the volumetric centroid. (B) Calculation for the magnitude of moment that is neglected by moving the force to the volumetric center and not including the moment in the resulting analysis.

### 2.3 Injury Simulation

In addition to modeling a SSM in an intact state, the model simulated the SSM in three levels of SLIL intrinsic ligamentous injuries^38^ by creating complete failures (removal of ligament from analysis) of all fibers of the affected ligaments at the start of the analysis^11^. These included isolated injuries to the VSL (“V”); combined VSL, and PSL (“VP”); and combined VSL, PSL, and DSL (“VPD”). No models contained injury to the extrinsic stabilizes. All other ligaments beyond VSL, PSL, and DSL, were left intact.

### 2.4 Output Metric Comparison

Output metrics included scaphoid to radius kinematics, radioscaphoid contact area (total area of cartilage with non-zero pressures), radioscaphoid contact force (net force magnitude), and average radioscaphoid contact pressure (force over total area), and the forces in each of the ligaments. Values were reported both as individual values within the motion cycle, and averages over the full motion cycle.

Model predictions were indirectly validated by comparing model (intact and VPD) predictions against previous work by Wolfe et al.^35^. Their study fluoroscopically measured the relationship between scaphoid flexion/extension and the dorsal translation in a SSM, with and without the dorsal scaphoid pressure applied. This work simulated this by creating an additional set of models performing the same SSM motion but without a dorsal scaphoid force. The differences in scaphoid rotation and dorsal translation were calculated between the models with and without the dorsal scaphoid force. As reported by others^35^, the dorsal translations were normalized to the total volar-dorsal distance of the distal radius and reported in “radius units” (RU). The points reported^35^ were digitized and overlaid against the values herein. Notably, in this work the curves created show the translation and flexion over the full motion cycle, whereas the original study reported only the difference at the final shift.

## 3 Results

In the intact wrist (Intact model), there was relatively constant contact pressure between the scaphoid and radius. In contrast, the model with a complete SLIL injury (VPD model) resulted in dorsal subluxation of the scaphoid at approximately 80% of the cycle, representing approximately 0° radioulnar deviation and 10° flexion, with a noticeable spike in contact area, force, and pressure (Figure 4). The motion of scaphoid in the SSM in the VPD model is qualitatively similar to fluoroscopically-captured SSM presented in Lui et al.^36^. The contact pressure region for the scaphoid on the radius in the Intact model was qualitatively similar across the motion cycle compared to the VPD model, where it began more volarly and moves dorsally over the motion cycle (Figure 4). Quantitatively, the scapholunate gap increased from 1.3 mm in the Intact model at 0% of the motion cycle to 1.8 mm in the VPD model.

**Figure 4.**
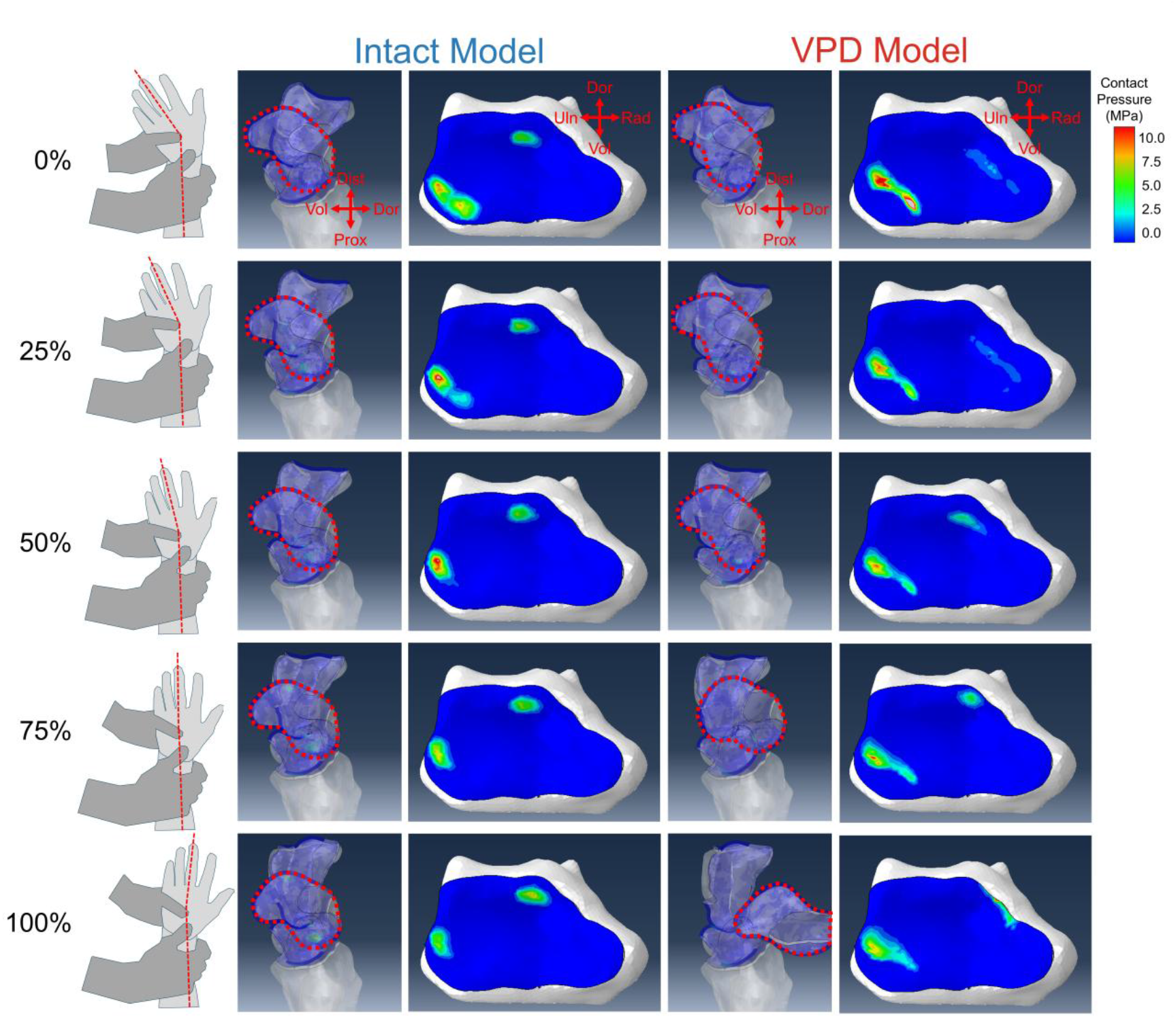
Simulated scaphoid shift maneuver (SSM) motion for the intact (intact SLIL) versus VPD (VSL, PSL, and DSL injury) models. Models demonstrate contact pressure plots with the outline of the scaphoid highlighted in dashed red region along the radial-ulnar direction. In addition, plots are shown for the distal radius showing the contact pressure along the distal-proximal direction. In the 75% of the motion cycle images, the scaphoid in the VPD model is noticeably more rotated in the pronosupination direction and has had additional scaphoid flexion and dorsal translation compared to the intact model The contact pressure region for the scaphoid on the radius in the intact model is relatively consistent. In contrast, this contact region in VPD model begins more volarly and moves dorsally over the motion cycle. Subluxation of the scaphoid in the VPD model occurs between 75%-100% of the motion cycle (at approximately 80% of the cycle).

### 3.1 Kinematics

The average scaphoid dorsal translation, relative to the intact model, was the largest just before the subluxation for the VPD model of 4.6 mm, followed by the V model with a dorsal shift of 3.0 mm (Figure 5). With injury, the scaphoid radial translation increased with injury up to an average of 1.3 mm in the VPD model relative to the intact model. Rotations were more variable than translations. Total scaphoid flexion across the motion cycle was largest in the VPD model at 24.6 degrees prior to subluxation. The pronosupination of the scaphoid had the largest joint angle range between models. The scaphoid was more radially deviated on average in all injury models compared to the intact model.

**Figure 5.**
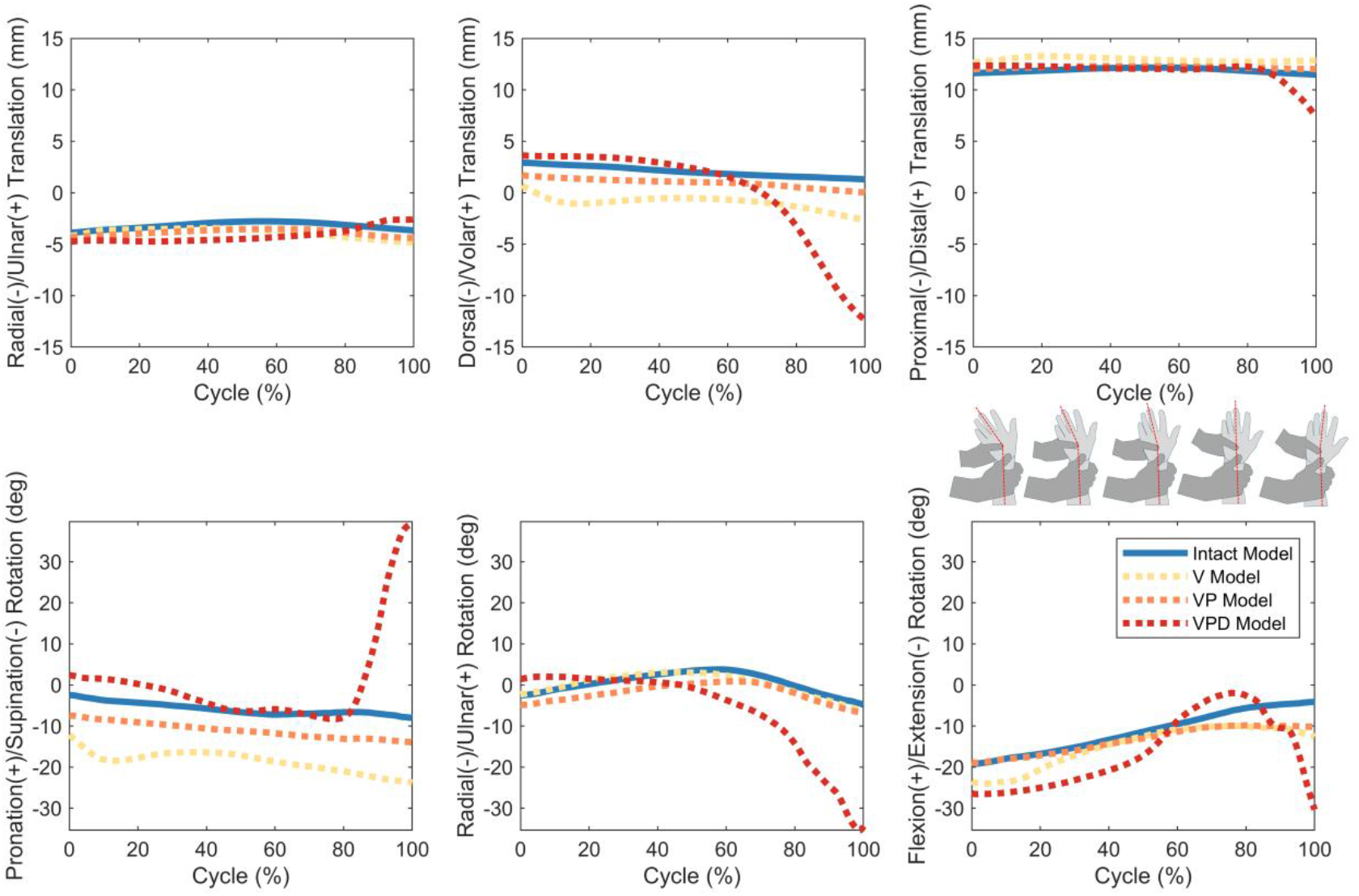
Six degree-of-freedom scaphoid kinematics relative to radius during the scaphoid shift maneuver (SSM) for the intact model (intact ligaments), V model (VSL injury), VP model (VSL and PSL injury), and the VPD model (VSL, PSL, and DSL injury). Subluxation of the scaphoid in the VPD model occurs at approximately 80% of the motion cycle.

Differences in scaphoid flexion and dorsal translation between the unloaded and loaded SSM were compared to the results presented by others^35^ and showed similar trends, namely that increased scaphoid flexion was associated with dorsal translation (Figure 6), particularly for the final motion cycle point.

**Figure 6.**
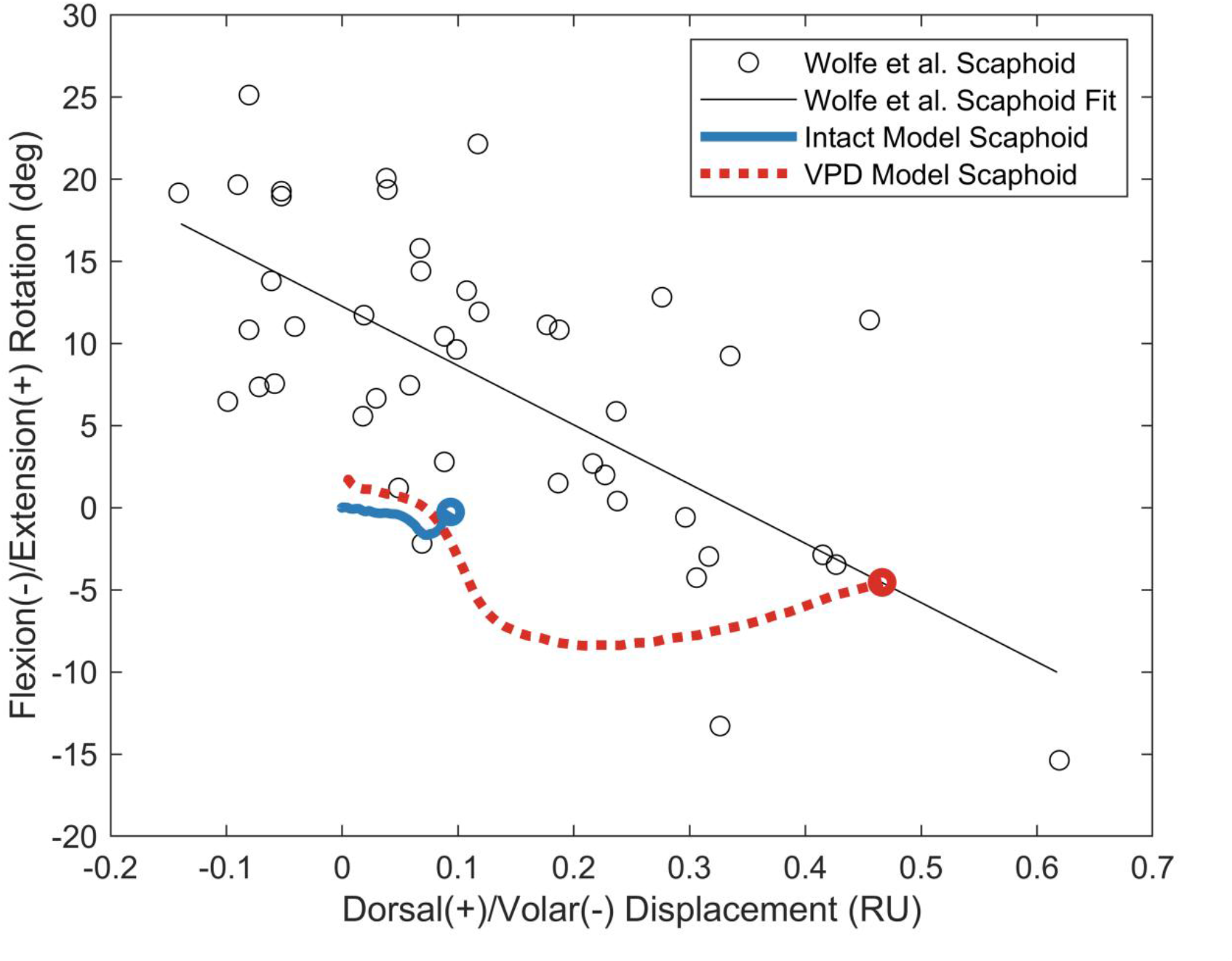
Recreation of dorsal displacement versus flexion/extension rotation plot from Wolfe et al. ^35^of the scaphoid during the scaphoid shift maneuver (SSM), with results from the intact (intact SLIL) and VPD (VSL, PSL, and DSL injury) models from the current study. Translations have been normalized to radial units (RU) by dividing the translation by the approximate total dorsal/volar distance of the distal radius. The VPD injury model has been truncated to the portion prior to scaphoid subluxation (0%-80% of motion cycle). Lines represent the complete SSM range of motion, while points from the prior work^35^ represent the static differences from individual participants. The corresponding points in this work (100% of the motion cycle for the intact, and instance of subluxation for VPD model) has been marked with a circle marker to enhance the comparison.

### 3.2 Contact Mechanics

At 0% of the motion cycle, the contact area was approximately 200% greater in the VPD model (Figure 7) compared with the contact area for the other models. Contact force increased for the intact, V, and VP models throughout the cycle, but decreased for the VPD model before spiking just before the scaphoid subluxation event. Contact pressure for all models increased as the cycle progressed. Except for the VPD model contact pressure just prior to subluxation, the V model had the highest cycle-average contact pressure.

**Figure 7.**
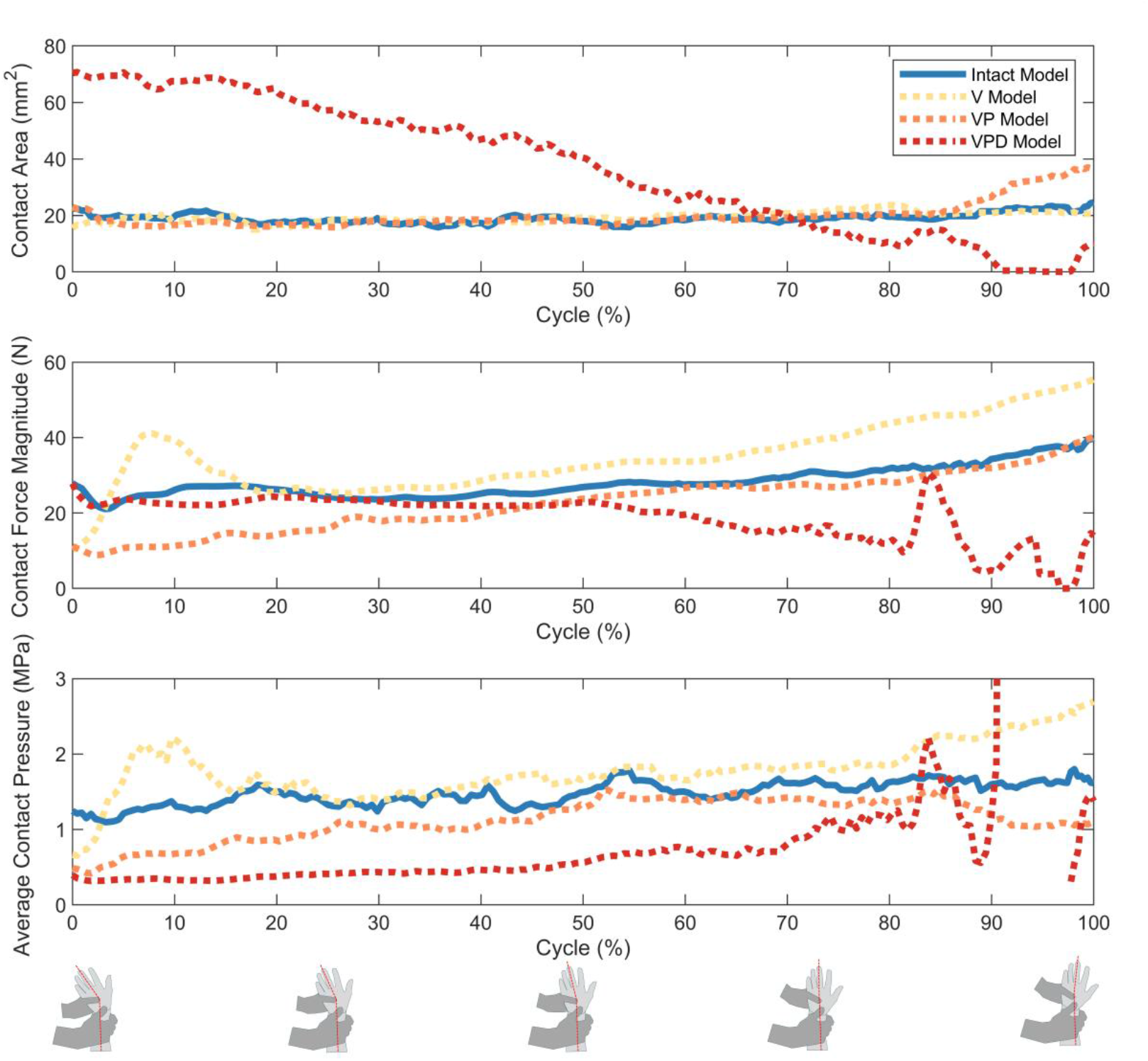
Scaphoid on radius cartilage contact area, contact force magnitude, and average contact pressure during the scaphoid shift maneuver (SSM) for the intact model (intact SLIL), V model (VSL injury), VP model (VSL and PSL injury), and the VPD model (VSL, PSL, and DSL injury). Note: The VPD model resulted in a subluxation (motion cycle ∼= 80%) that yielded very high contact pressures, up to 80 MPa, driven by the very small contact areas. To optimize visualization of the other lines, the y-axis has been truncated to a maximum pressure of 3 MPa, as values past this exist only for the VPD model following subluxation, which computationally are prone to significant errors and often physically meaningless.

### 3.3 Ligament Forces

Ligament forces changes (relative to the intact model) varied greatly but were greatest in the VPD model, followed by the V model, and then the VP model relative to the intact model. Forces in the UCL, UL, and RCL were less than 5.0 N through the entire motion cycle in all models. For the SLIL, the forces in the PSL and VSL differed more than the forces in the DSL between injury models across the motion cycle (Figure 8). The force in the VSL (33.0 N maximum, 14.4 N average) was lower than the DSL (54.2 N, 22.6 N average) in the intact model; however, differences were not constant throughout the motion cycle. The force in the PSL was less than 1.0 N for all models except for the V model, where force increased (17.9 N maximum, 10.5 N average). The LRL, an extrinsic stabilizer of the scapholunate joint, had forces that were greater in the VP and VPD models compared to the intact model. In contrast, the ligament forces in the SCL, not usually considered a scapholunate stabilizer, decreased in the V, VP, and VPD models. Ligament force changes in the SRL, RSC, and DRC were observed but no consistent trends were found.

**Figure 8.**
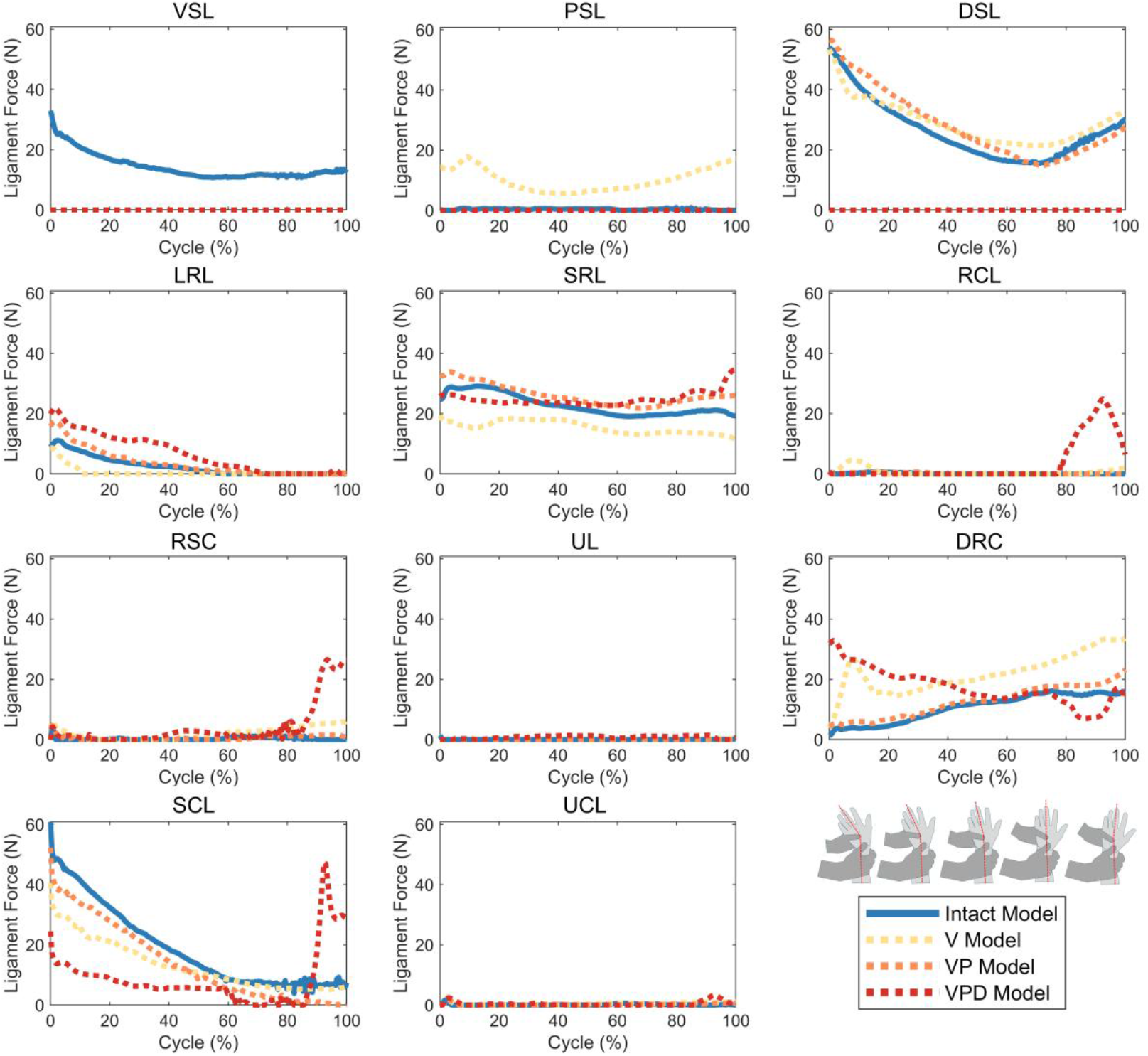
Ligament forces in modeled ligaments during the scaphoid shift maneuver (SSM) for the intact model (intact SLIL), V model (VSL injury), VP model (VSL and PSL injury), and the VPD model (VSL, PSL, and DSL injury). Subluxation of the scaphoid in the VPD model occurs at approximately 80% of the motion cycle.

## 4 Discussion

Using a personalized wrist model this work simulated an SSM under different SLIL injuries (Figure 1). The SSM was chosen to represent a clinically-relevant provocative maneuver. The complete intrinsic SLIL injury model (VPD model) captured the rapid dorsal shift and subluxation of the scaphoid (Figure 4 and Figure 5) associated with a positive SSM^14,15^, as well as the radially translated scaphoid and increased scapholunate gap frequently associated with SLIL injuries^20,22^. In addition, difference between the unloaded and loaded SSM (Figure 6) yielded a similar relationship between dorsal displacement and flexion angle to that reported by others ^35^. The ability of the models to recreate the complex dynamics of an SSM, including scaphoid subluxation, demonstrates the usefulness of our modeling approach over existing work that has been largely limited to static analyses^12^.

This work demonstrates how these models can offer new insights via quantification of metrics that would be otherwise difficult to measure. For example, in the V model, the contact force and average contact pressure increased relative to the fully intact model (Figure 7). Increased joint contact pressure has previously been shown to be a predictor of osteoarthritis (OA) in other joints^39^ and may explain the progression of untreated SLIL injuries to scapholunate advanced collapse pattern OA^19^. Still, changes in average contact pressure alone (total force over total contact area) may not be the most appropriate indicators of wrist OA progression^11^; rather changes in the location of contact pressure may be just as important and could be the focus of future studies.

The ligament force changes observed may explain the progression of SLIL ligament injuries (Figure 8). The maximum ligament forces in the intact wrist during the SSM was 33.0 N and 54.2 N for the VSL and DSL, respectively. Previous testing shows the rupture strengths to be 117.9 N 260.3 N for the VSL and DSL, respectively ^40^. As a proportion of its overall strength (force over strength), the forces in the VSL are higher than the DSL and may explain why it is frequently injured first^21^. Similarly, the increased force sharing of the PSL following VSL injury (maximum force of 17.9N for a rupture strength of 62.7) may explain why the next ligament injured is frequently the PSL compared with the DSL^21^. The relatively consistent force in the DSL across injury states, combined with increasing forces in the LRL for more severe injuries, provides evidence for the repair of extrinsic stabilizers.

This work is not free of limitations. This work used an anatomical model derived from one asymptomatic participant. The models predicted subluxation of the scaphoid only in the most severe VPD injury case, suggesting that the model can accurately recreate aspects of intact joint dynamics. When these dynamics are altered, they reflect the behaviors of the injured state observed clinically. This work demonstrates how these types of models may be used to simulate not only common biomechanical tests but also important clinical tests. Still, the small sample size inherently limits the broad generalization of claims. Future work should aim to include a greater number of participants, including those with arthroscopically confirmed injuries. Additionally, further simulations should be performed to determine if the observed patterns persist, particularly in more complex injury conditions and across broader patient demographics. The other limitation is the lack of an anatomically complete wrist model. While this work was able to show the initial subluxation, it was unable to replicate the “clunk”, that is often clinically detected, as no structures were modeled that would return the scaphoid to its initial position following the removal of the applied dorsal force. The inclusion of additional biological structures may better recreate this phenomenon. Still, the structures modeled match those of prior work^41^, and as shown, can recreate the key aspects of the SSM: rapid dorsal translation and flexion of the scaphoid and, in extreme cases, subluxation.

In conclusion, this work used a personalized computational model of an asymptomatic participant to simulate the effects of a SSM at different SLIL injury levels. Model outcomes highlighted differences in kinematics, contact mechanics, and ligament forces between injuries. Models were able to recreate dynamic behavior of the injury, particularly demonstrating scaphoid subluxation in the fully injured model. The models herein captured contact mechanics changes that may be important for studying the interplay between soft-tissue injury and long-term joint mechanics that may be related to OA development. Lastly, this work identified changes to ligament forces that may explain the progression of injury from VSL to complete intrinsic SLIL injury, to the involvement of extrinsic ligaments. This work enables future investigations into how SLIL injuries may impact patients both acutely and chronically.

## 7 Acknowledgements

The authors would also like to thank Altair Engineering Inc. for generously providing HyperWorks software to enable this work. The authors would also like to thank the Mayo Clinic Computed Tomography Clinical Innovation Center for invaluable contributions to data collection as well as the Bio-Imaging Research Core at Mayo Clinic for their help with this work.

## References

1. Innocenti B, Bori E, Armaroli F, et al. The use of computational models in orthopedic biomechanical research. Human Orthopaedic Biomechanics. 2022:681–712.

2. Imhauser CW, Baumann AP, Liu XC, et al. Reproducibility in Modeling and Simulation of the Knee: Academic, Industry, and Regulatory Perspectives. Journal of Orthopaedic Research 2023.

3. Goetz LH, Schork NJ. Personalized medicine: motivation, challenges, and progress. Fertil Steril. Jun 2018;109(6):952–963. doi:10.1016/j.fertnstert.2018.05.006

4. Sun T, He X, Song X, Shu L, Li Z. The Digital Twin in Medicine: A Key to the Future of Healthcare? Frontiers in Medicine 2022. p. 1–8.

5. Sun T, Wang J, Suo M, et al. The Digital Twin: A Potential Solution for the Personalized Diagnosis and Treatment of Musculoskeletal System Diseases. Bioengineering 2023. p. 627.

6. Viceconti M, De Vos M, Mellone S, Geris L. Position paper From the digital twins in healthcare to the Virtual Human Twin: a moon-shot project for digital health research. IEEE J Biomed Health Inform. Oct 11 2023;PP doi:10.1109/JBHI.2023.3323688

7. Pathmanathan P, Aycock K, Badal A, et al. Credibility assessment of in silico clinical trials for medical devices. PLoS Comput Biol. Aug 2024;20(8):e1012289. doi:10.1371/journal.pcbi.1012289

8. Andres A, Roland M, Wickert K, et al. Advantages of digital twin technology in orthopedic trauma Surgery - Exploring different clinical use cases. Sci Rep. Jun 6 2025;15(1):19987. doi:10.1038/s41598-025-04792-w

9. Maquer G, Mueri C, Henderson A, Bischoff J, Favre P. Developing and Validating a Model of Humeral Stem Primary Stability, Intended for In Silico Clinical Trials. Ann Biomed Eng. May 2024;52(5):1280–1296. doi:10.1007/s10439-024-03452-w

10. Delport H, Mulier M, Gelaude F, Clijmans T. Complex Acetabular Revision Using Computer-Aided Planning for Patient-Specific Implant and Guide. Orthopaedic Proceedings. 2012;94 doi:10.1302/1358-992X.94BSUPP_XXV.ISTA2010-040

11. Andreassen TE, Trentadue TP, Thoreson AR, An KN, Kakar S, Zhao KD. Rapid Development of Efficient Participant-Specific Computational Models of the Wrist. Preprint. 2025;doi:10.48550/arXiv.2505.19282

12. Mena A, Wollstein R, Baus J, Yang J. Finite Element Modeling of the Human Wrist: A Review. J Wrist Surg. Dec 2023;12(6):478–487. doi:10.1055/s-0043-1768930

13. Alonso Rasgado T, Zhang Q, Jimenez Cruz D, et al. Analysis of tenodesis techniques for treatment of scapholunate instability using the finite element method. Int J Numer Method Biomed Eng. Dec 2017;33(12)doi:10.1002/cnm.2897

14. Watson HK, Ashmead Dt, Makhlouf MV. Examination of the scaphoid. J Hand Surg Am. Sep 1988;13(5):657–60. doi:10.1016/s0363-5023(88)80118-7

15. Watson HK, Ryu J, Akelman E. Limited triscaphoid intercarpal arthrodesis for rotatory subluxation of the scaphoid. J Bone Joint Surg Am. Mar 1986;68(3)(3):345-9.

16. Schmauss D, Pohlmann S, Weinzierl A, et al. Relevance of the Scaphoid Shift Test for the Investigation of Scapholunate Ligament Injuries. J Clin Med. Oct 26 2022;11(21)doi:10.3390/jcm11216322

17. Prosser R, Harvey L, Lastayo P, Hargreaves I, Scougall P, Herbert RD. Provocative wrist tests and MRI are of limited diagnostic value for suspected wrist ligament injuries: a cross-sectional study. J Physiother. 2011;57(4):247–53. doi:10.1016/S1836-9553(11)70055-8

18. Vilai P, Kakar S. The Diagnostic Value of the Scapholunate C-Sign: A New Tool for Detecting Through-and-Through Scapholunate Interosseous Ligament Injuries. J Hand Surg Am. Aug 14 2025;doi:10.1016/j.jhsa.2025.06.011

19. Watson HK, Ballet FL. The SLAC wrist: scapholunate advanced collapse pattern of degenerative arthritis. J Hand Surg Am. May 1984;9(3):358–65. doi:10.1016/s0363-5023(84)80223-3

20. Zhou JY, Jodah R, Joseph LP, Yao J. Scapholunate Ligament Injuries. J Hand Surg Glob Online. May 2024;6(3):245–267. doi:10.1016/j.jhsg.2024.01.015

21. Garcia-Elias M, Lluch AL, Stanley JK. Three-ligament tenodesis for the treatment of scapholunate dissociation: indications and surgical technique. J Hand Surg Am. Jan 2006;31(1):125–34. doi:10.1016/j.jhsa.2005.10.011

22. Geissler WB. Arthroscopic management of scapholunate instability. J Wrist Surg. May 2013;2(2):129–35. doi:10.1055/s-0033-1343354

23. Trentadue TP, Thoreson A, Lopez C, et al. Morphology of the scaphotrapeziotrapezoid joint: A multi-domain statistical shape modeling approach. J Orthop Res. Nov 2024;42(11):2562–2574. doi:10.1002/jor.25918

24. Trentadue TP, Thoreson A, Lopez C, et al. Sex differences in photon-counting detector computed tomography-derived scaphotrapeziotrapezoid joint morphometrics. Skeletal Radiol. Feb 5 2025;doi:10.1007/s00256-024-04863-5

25. Trentadue TP, Thoreson AR, Lopez C, et al. Detection of scapholunate interosseous ligament injury using dynamic computed tomography-derived arthrokinematics: A prospective clinical trial. Med Eng Phys. Jun 2024;128:104172. doi:10.1016/j.medengphy.2024.104172

26. Trentadue TP, Lopez C, Thoreson A, et al. Four-dimensional computed tomography-derived distal radioulnar joint osteokinematics during pronosupination with unilateral triangular fibrocartilage complex injuries. J Biomech. Jul 2025;188:112703. doi:10.1016/j.jbiomech.2025.112703

27. Trentadue TP, Lopez C, Breighner RE, et al. Evaluation of Scapholunate Injury and Repair with Dynamic (4D) CT: A Preliminary Report of Two Cases. J Wrist Surg. Jun 2023;12(3):248–260. doi:10.1055/s-0042-1758159

28. Zhao K, Breighner R, Holmes D, 3rd, Leng S, McCollough C, An KN. A technique for quantifying wrist motion using four-dimensional computed tomography: approach and validation. J Biomech Eng. Jul 2015;137(7):0745011–5. doi:10.1115/1.4030405

29. Andreassen TE, Hume DR, Hamilton LD, Higinbotham SE, Shelburne KB. Automated 2D and 3D finite element overclosure adjustment and mesh morphing using generalized regression neural networks. Medical Engineering & Physics. 2024; 126 doi:10.1016/j.medengphy.2024.104136

30. Andreassen TE, Hume DR, Hamilton LD, Hegg SL, Higinbotham SE, Shelburne KB. Validating subject-specific knee models from in vivo measurements. Frontiers in Bioengineering and Biotechnology. 2025;13(1554836)doi:10.3389/fbioe.2025.1554836

31. Hume DR, Rullkoetter PJ, Shelburne KB. ReadySim: A computational framework for building explicit finite element musculoskeletal simulations directly from motion laboratory data. International Journal for Numerical Methods in Biomedical Engineering 2020. p. 1–11.

32. ASME. Assessing Credibility of Computational Modeling through Verification and Validation: Application to Medical Devices. V&V40: ASME; 2018. p. 1–48.

33. Coburn JC, Upal MA, Crisco JJ. Coordinate systems for the carpal bones of the wrist. J Biomech. 2007;40(1):203–209. doi:10.1016/j.jbiomech.2005.11.015

34. Neu CP, Crisco JJ, Wolfe SW. In vivo kinematic behavior of the radio-capitate joint during wrist flexion-extension and radio-ulnar deviation. J Biomech. Nov 2001;34(11):1429–38. doi:10.1016/s0021-9290(01)00117-8

35. Wolfe SW, Gupta A, Crisco JJ, III. Kinematics of the scaphoid shift test. J Hand Surg Am. Sep 1997;22(5):801–6. doi:10.1016/S0363-5023(97)80072-X

36. Lui TH, Slocum AMY. Scaphoid Shift Test in Scapholunate Ligament Injury. N Engl J Med. Nov 3 2022;387(18):e46. doi:10.1056/NEJMicm2202238

37. Wolfe SW, Crisco JJ. Mechanical evaluation of the scaphoid shift test. J Hand Surg Am. Sep 1994;19(5):762–8. doi:10.1016/0363-5023(94)90180-5

38. Trentadue TP, Lopez C, Breighner RE, et al. Assessing carpal kinematics following scapholunate interosseous ligament injury ex vivo using four-dimensional dynamic computed tomography. Clin Biomech (Bristol). Jul 2023;107:106007. doi:10.1016/j.clinbiomech.2023.106007

39. Segal NA, Anderson DD, Iyer KS, et al. Baseline articular contact stress levels predict incident symptomatic knee osteoarthritis development in the MOST cohort. J Orthop Res. Dec 2009;27(12):1562–8. doi:10.1002/jor.20936

40. Berger RA, Imeada T, Berglund L, An KN. Constraint and material properties of the subregions of the scapholunate interosseous ligament. J Hand Surg Am. Sep 1999;24(5):953–62. doi:10.1053/jhsu.1999.0953

41. Marques R, Melchor J, Sanchez-Montesinos I, Roda O, Rus G, Hernandez-Cortes P. Biomechanical Finite Element Method Model of the Proximal Carpal Row and Experimental Validation. Front Physiol. 2021;12:749372. doi:10.3389/fphys.2021.749372

